# Variation in traction forces during cell cycle progression

**DOI:** 10.1101/212472

**Authors:** Benoit Vianay, Fabrice Senger, Simon Alamos, Maya Anjur-Dietrich, Elizabeth Bearce, Bevan Cheeseman, Lisa Lee, Manuel Théry

## Abstract

Tissue morphogenesis results from the interplay between cell growth and mechanical forces. While the impact of forces on cell proliferation has been fairly well characterized, the inverse relationship is much less understood. Here we investigated how traction forces vary during cell cycle progression. Cell shape was constrained on micropatterned substrates in order to distinguish variations in cell contractility from cell size increase. We performed traction force measurements of asynchronously dividing cells expressing a cell-cycle reporter, to obtain measurements of contractile forces generated during cell division. We found that forces tend to increase as cells progress through G1, before reaching a plateau in S phase, and then decline during G2. This biphasic behaviour revealed a previously undocumented specific and opposite regulation of cell contractility during each cell cycle stage.

## Introduction

Tissue morphogenesis, during both embryo development and adult tissue renewal, relies on cell growth and shape changes (Thompson, 1942; Lecuit and Lenne, 2007). Tissue growth is mostly supported by cell proliferation. The determination of tissue shape depends on the production of mechanical forces that regulate cell morphology and position (Heisenberg and Bellaïche, 2013). Tissue shape also depends on the spatial regulation of cell differentiation (Heller and Fuchs, 2015; Maitre *et al.*, 2016; Gilmour *et al.*, 2017). Cell mechanics, fate, and growth are far from independent, and the spatio-temporal coordination of growth, differentiation and shape acquisition relies on a tight coupling between the three. It is widely-established that mechanical forces and cell shape direct cell fate and regulate cell cycle progression (Watt *et al.*, 1988; Chen *et al.*, 1997; Ruiz and Chen, 2008; Guilak *et al.*, 2009; Klein *et al.*, 2009; Kilian *et al.*, 2010; Dupont *et al.*, 2011; Chan *et al.*, 2017).

The impact of mechanical forces on cell growth has been the focus of numerous studies, but much less is known about causality in the opposite direction; i.e the effect of cell cycle progression on the production of mechanical forces. Growth factor starvation showed that quiescent cells produce less force than proliferating cells (Rape *et al.*, 2011b). The dynamics of mechanical forces produced across the cell cycle are largely unknown, though studies have nicely-characterized aspects of force production explicitly during mitosis. As cells enter mitosis, they detach from the extra-cellular matrix in a process called deadhesion (Marchesi *et al.*, 2014) resulting in a drastic reduction of tractional forces (Lesman *et al.*, 2014). Mitotic cells continue to produce contractile forces, but they are distributed internally and lead to cell rounding and stiffening (Maddox and Burridge, 2003; Théry and Bornens, 2008). Cells regain the ability to produce traction forces as they exit from mitosis and respread onto the extra-cellular matrix in early G1 (Cramer and Mitchison, 1995; Lesman *et al.*, 2014).

It is not known how traction forces vary from early G1 to late G2. The null hypothesis is that they remain constant, however, the main characteristic of cell cycle progression is cell growth: cell size and mass increase steadily from early G1 to late G2 (Kafri *et al.*, 2013; Son *et al.*, 2015; Varsano *et al.*, 2017). Several works have shown that cell size has a clear influence on the production of traction forces, and that bigger cells tend to produce larger forces (Tan *et al.*, 2003; Reinhart-king *et al.*, 2005; Tolić-Nørrelykke and Wang, 2005; Rape *et al.*, 2011a; Oakes *et al.*, 2014). According to this trend, traction forces should increase steadily with cell cycle progression. We took advantage of a two-week rotation during the Physiology course in Woods Hole to test these hypotheses, and measure the evolution of traction forces during cell cycle progression.

## Results and Discussion

One straightforward strategy to assess traction forces across the cell cycle would rely on synchronizing cells and performing force production measurements during each cell cycle stage. However synchronizing drugs, which inhibit specific cyclin kinases, blocks DNA replication or disassemble microtubules (Ma and Poon, 2017), can interfere with normal cell cycle progression after release (Bar-Joseph *et al.*, 2008). Rather than pharmacologically perturbing the cell cycle to induce synchronization, we opted to utilize asynchronous cells expressing the fluorescent ubiquitin-based cell cycle indicator (FUCCI) reporter system. The FUCCI reporter is based on the sequential hCdt1-mCherry expression in G1 and hGem-Azami Green expression in S/G2/M (Sakaue-Sawano *et al.*, 2008). We worked with RPE-1 cells, a diploid, nontransformed human epithelial cell line, stably expressing the Fucci constructs (Ganem *et al.*, 2014) (Figure 1A). Cells were plated on soft poly-acrylamide gel with embedded fiduciary beads, to visualize gel deformation and infer the traction forces produced by the cells, as previously described (Dembo and Wang, 1999) (Figure 1B). It is important to plate cells at low density in order to detect their individual traction force field. However, RPE1 cells are motile in these conditions, and migration is a great source of variability in force production (Meili *et al.*, 2010; Chang *et al.*, 2013; Leal-Egaña *et al.*, 2017). In order to limit these large variations that could blur the changes due to cell cycle progression, cells were plated on adhesive micropatterns, which prevented their motion and normalized their morphology, to achieve a constant and reproducible shape (Singhvi *et al.*, 1994; Théry, 2010). We further considered that standardizing stress fiber position and number would reduce inter-cellular variability (Mandal *et al.*, 2014). We achieved this by plating cells on 60-micron-long and 12-micro-wide dumbell-shaped micropatterns, in which the shape and position of non-adhesive regions dictate the number, size and position of stress fibers (Théry *et al.*, 2006) (Figure C). The combination of these methods: the Fucci reporter, the deformable substrate and the controlled cell shape, allowed us to measure cell cycle position and traction forces in standardized conditions (Figure 1D).

**Figure 1:**
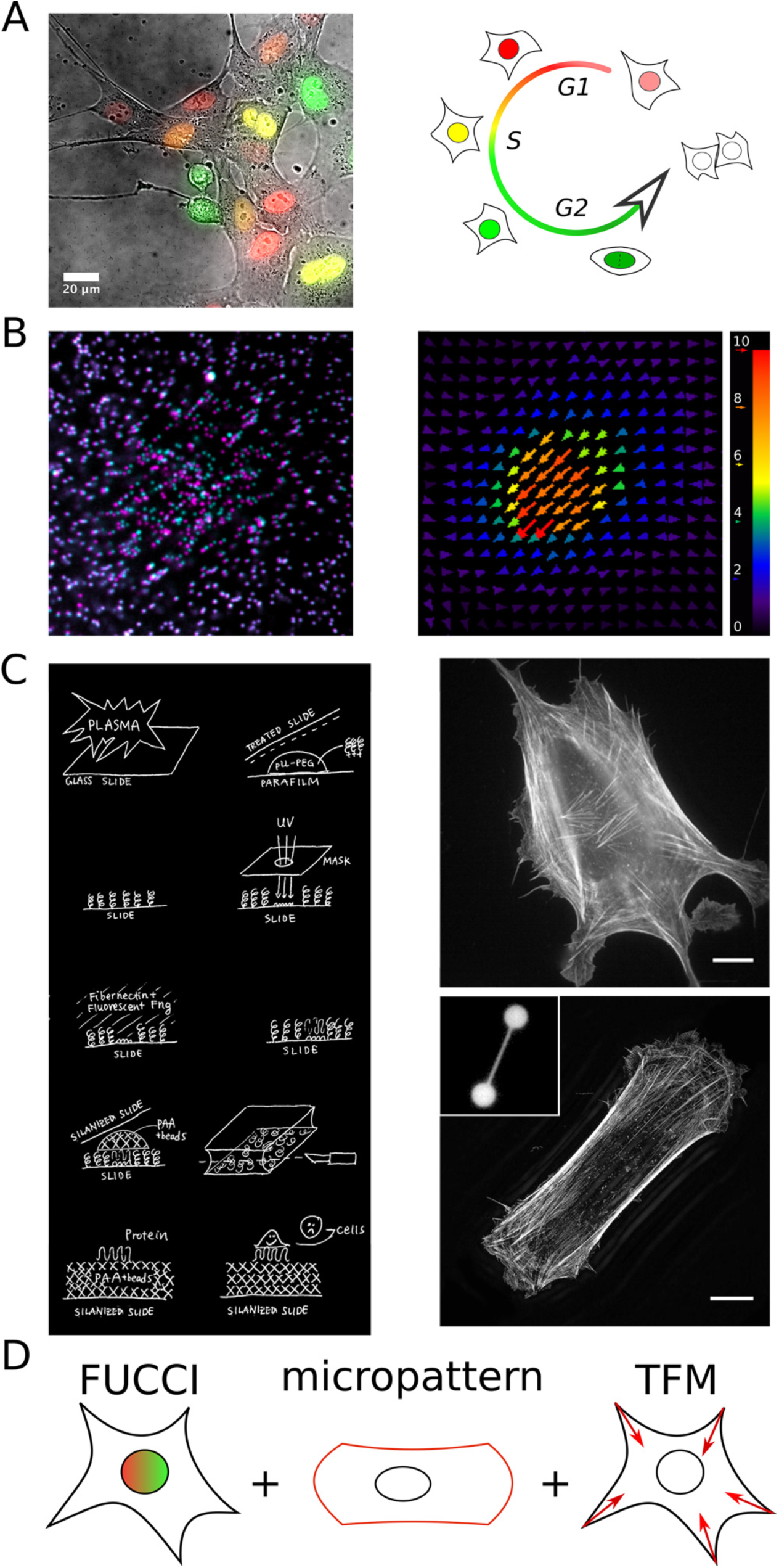
Experimental set up. A)Left: RPE1 FUCCI cells spread on PAA gels homogeneously coated with fibronectin. The color of the nucleus indicates its cycle phase. Right: Nucleus color changes during cell cycle progression. B)Left: Typical images of the beads displacements due to cell traction forces (magenta) compared to beads with no stress (cyan). Right: PIV results of the image on the left with the corresponding beads displacement field. Displacement scale bar is in pixels. C)Left: Procedure to fabricate micropatterns on PAA gels. 1st row: glass pegylation after plasma activation; 2nd row: DeepUV illumination; 3rd row: protein incubation; 4th row: PAA gel polymerisation and separation; 5th row: resulting protein micropatterns on PAA gel and cell seeding (see Material and Methods). Right: RPE1 cell spread on homogeneous fibronectin coated glass substrate (top), or on fibronectin patterned glass substrate (bottom) where actin stress fibers are well defined at the two edges due to dumbbell pattern (inset). These two images were acquired by the DeltaVision OMX SR (GE Healthcare). D)Experimental set up based on FUCCI cells spread on micro patterned PAA gels to measure cell traction forces during cell cycle progression.

We first confirmed that cells displayed the expected color changes as they progressed in the cell cycle when micropatterned on poly-acrylamide gel (Figure 2). Fibronectin-coated micropatterns were first manufactured on glass coverslips, and then transferred onto a poly-acrylamide hydrogel (Vignaud *et al.*, 2014) (see Methods). RPE1-Fucci cells were plated on micropatterned gels and monitored 24h using time-lapse confocal microscopy. As expected, cells expressing exclusively the hCdt1-mCherry (red) construct at the beginning of the cell cycle, reduced it progressively over time, and increased the production of hGem-Azami Green, resulting in the exclusive production of hGem-Azami Green approximately 12hrs later, at the end of S phase (Figure 2A). This « green » phase, which corresponded to the G2 phase, lasted about five hours until entry into mitosis (Figure 2A).

**Figure 2:**
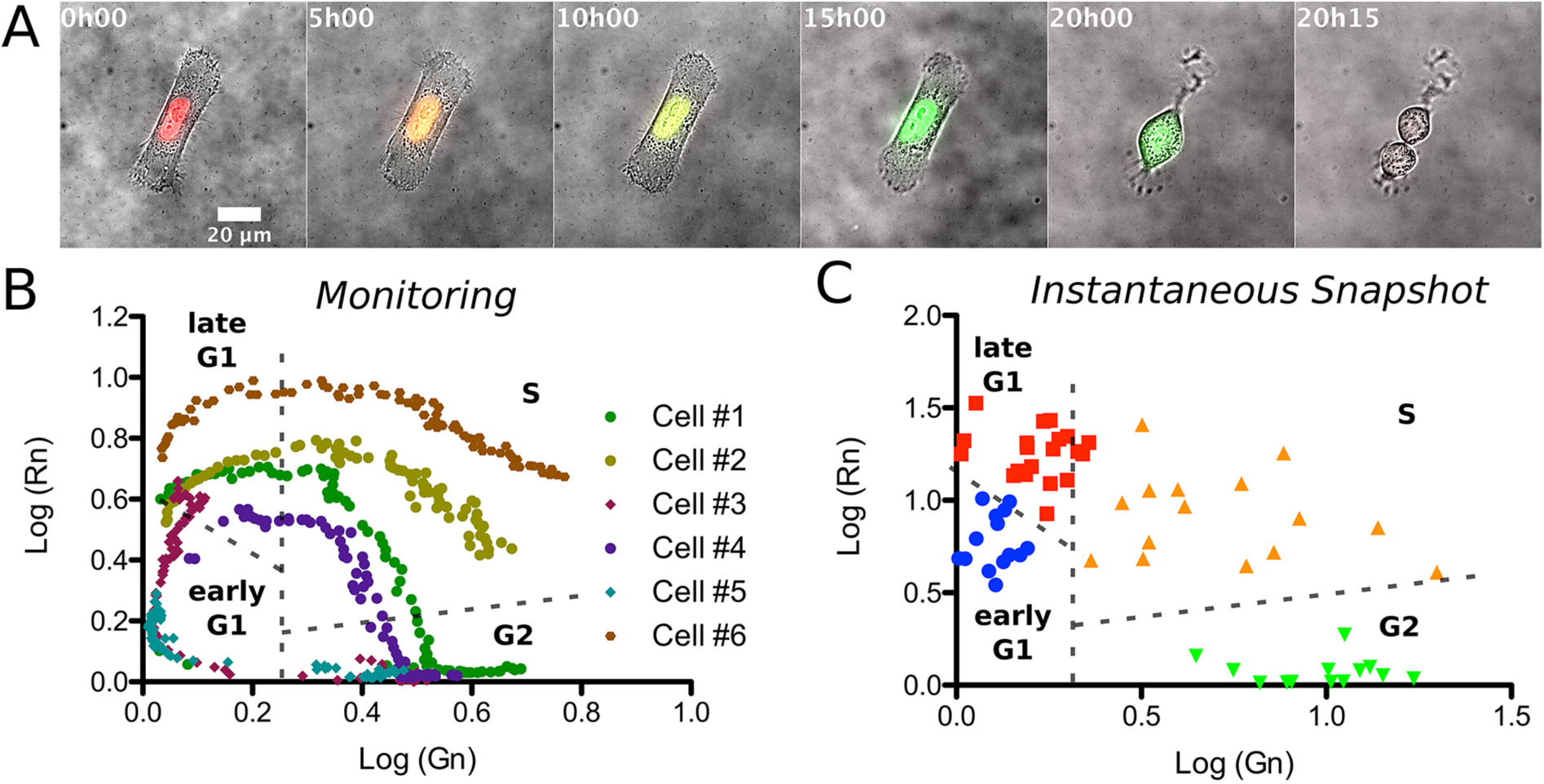
Determining cell cycle progression. A)Timelapse phase with red and green fluorescence combined images of a RPE1 FUCCI cell spread on a dumbbell pattern on PAA gels over 20 hours. The nucleus color changes confirm the cell cycle progression from G1 - S - G2 until mitosis at 20h15 after monitoring. B)Log-log diagram of the nuclei normalized red color in function of the nuclei normalized green color for 6 different cells monitored over 20 hours during cycle progression. The dashed lines are guides to separate cycle phases (early G1, late G1, S and G2 phases). Diamonds correspond to mitotic cells (G2 - M - G1) during monitoring whereas circles correspond to cells progressing from G1 to G2. The green circles (cell #1) corresponds to the cell in A). C)Same log-log diagram of the nuclei normalized color than in B), each point represent a cell (N=3 independant experiments). The colors correspond to the different cycle phases defined by the dashed lines in B).

These durations approximately correspond to the cell cycle durations reported for this cell line (Azimzadeh *et al.*, 2009). When the fluorescence signal of each reporter is plotted over time for individual cells, their trajectories follow the typical, dome-like, trend of normal cell cycle progression (Figure 2B, to be compared to Figure 1G in (Sakaue-Sawano *et al.*, 2008)). Similarly, the balance of fluorescence intensities in individual cells at a given time point displayed the same distribution (Figure 2C). Previous characterization of the relationship between fluorescence ratio and cell cycle stage (Sakaue-Sawano *et al.*, 2008) were used to define the boundaries separating the various cell cycle stages (Figure 2B,C). Unfortunately very few cells were detected in the earliest phase of G1 (corresponding to the bottom left corner of the graph, ie absence of hGem-Azami Green and low level of hCdt1-mCherry) since the cell detachment, plating and spreading processes took several hours.

Once the cell cycle stage had been determined by measuring the fluorescence intensities of the two reporters, we imaged the dark-red-fluorescent beads that were embedded in the poly-acrylamide gel, to obtain their position while under tension. Cells were then treated with trypsin to disengage the traction forces that were applied on the substrate, and allow relaxation of the fiducial beads. Images of the beads in the presence and absence of cell-mediated tension were processed in order to measure their auto-correlation function and deduce the gel deformation field (Tseng *et al.*, 2012; Martiel *et al.*, 2015) (see Methods). We then used Fourier-transform traction cytometry to estimate the corresponding cell traction force field (Butler *et al.*, 2002; Martiel *et al.*, 2015). (Figure 3A). The force field was further used to calculate the total traction energy produced by each individual cell, to generate the substrate deformation we observed (Butler *et al.*, 2002; Martiel *et al.*, 2015). We then combined the measure of cell cycle position (Figure 2C) and the values of traction energies for each individual cell, to plot the variations of traction forces with respect to cell cycle progression (Figure 3B). To that end, we used cell position along a curvilinear axis representing cell cycle progression in the Fucci reporter graph as a proxy for cycle state (Figure 3B). We observed a biphasic evolution of traction forces. Traction forces first increased from early G1 to late G1 and S phase (Figure 3B). More surprisingly, traction forces then dropped after S phase until G2 (Figure 3B). Cells were further classified with respect to their cycle stage based on the boundaries established previously (Sakaue-Sawano *et al.*, 2008), in order to compare the forces produced in the distinct cell cycle stages. We confirmed that cells entering S phase produced significantly higher traction forces than cells in early G1 or late G2 (Figure 3C).

**Figure 3:**
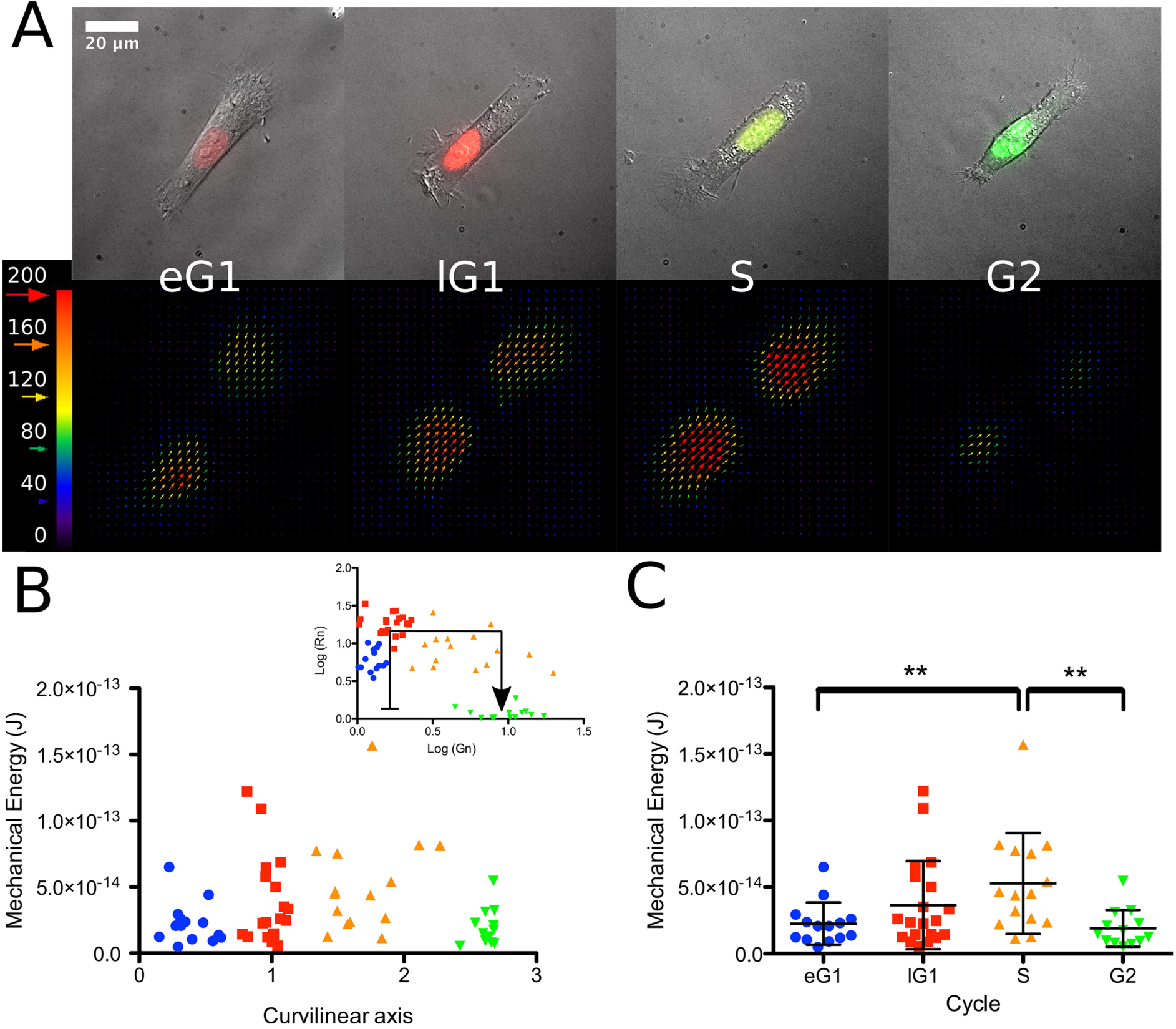
Traction forces variation during cell cycle progression. A)Phase with red and green fluorescence combined images (top row) of representative instantaneous snapshots of cells at early G1, late G1, S and G2 phase with their corresponding traction force fields (bottom row). For The grid size is 3.3 µm X 3.3 µm for a clear visualization of traction-force fields. Force scale bar is in Pascal. B)Cell mechanical energy along curvilinear axis (black arrow) defined in the log-log diagram of nuclei colors (inset). A clear increase of mechanical energy is observed during the S phase. C)Statistics of mechanical energy of early G1, late G1, S and G2 grouped cells extracted from B). Error bars represent mean values and standard deviations (** p<0.01).

The increase of forces from G1 to S is consistent with previous predictions relating cell area and contractility (Tan *et al.*, 2003; Reinhart-king *et al.*, 2005; Tolić-Nørrelykke and Wang, 2005; Rape *et al.*, 2011a; Oakes *et al.*, 2014), based on the fact that cell mass and volume increase from G1 to S phase (San JCB15, Son Varsano CellRep 17). However, in our experimental setting, cell area is predetermined by the micropatterned substrate; effectively uncoupling cell cycle progression from the contact area. Since all cells had the exact same spreading and adhesion areas, the increase in traction forces must reflect a genuine activation of the traction force machinery from early G1 to S phase. The force reduction after S phase was unexpected, given that cell mass and volume keep increasing during this period (Kafri *et al.*, 2013; Son *et al.*, 2015; Varsano *et al.*, 2017). These force variations may reflect changes in integrin activation. Indeed, integrins are specifically activated by growth factors during G1 (Walker and Assoian, 2005), so this phase may be more effective in force production. After the G1/S transition, cells are committed to mitosis and their progression is irreversible. In S and G2, cells are no longer sensitive to growth factors. The off-switching of their receptors is likely to impact the integrin activation and be responsible for the reduction in forces that we observed. The mechanism responsible for the force variations we observed deserves further investigation. Additionally, it is important to extend this study on single cells to the tissue level. How do intercellular tensional forces vary during cell cycle progression? How do cells sense the changes in traction and tension in their neighbours? Does it impact their own cell cycle progression in a global mechanical regulation of tissue homeostasis? These important questions will require more than a two-week practical course in Woods Hole to be addressed but may now be built on this primary study.

## Materials and Methods

### Cell lines

RPE1-FUCCI (provided by the lab of David Pellman) were grown in a humidified incubator at 37°C and 5% CO_2_ in DMEM/F12 medium supplemented with 10% fetal bovine serum and 1% penicillin/streptomycin. All cell culture products were purchased from GIBCO/Life technologies.

Cells were seeded on patterned gels at 100,000 cells/cm^2^. Non-adherent cells were washed away as soon as cells started to attach to the micropatterns. Cells were then allowed to spread fully onto the patterns for 3 hours.

### Hydrogel Micropatterning

Detailed procedure has been described elsewhere for glass micropatterning (Azioune *et al.*, 2010). and gel micropatterning (Vignaud *et al.*, 2014). In brief, glass coverslips were oxidized by oxygen plasma (PDC-100-HP Harrick Plasma) (10 sec, 30 W) and incubated for 30 min. with 0.1 mg/ml PLL-g-PEG (PLL20K-G35-PEG2K, JenKem) in 10mM HEPES pH 7.4. Dried coverslips where then exposed to deep-UV (PSD Pro series NOVASCAN) through a photomask (Toppan) for 4 min. After UV treatment, coverslips were incubated with 10 µg/ml fibronectin (Sigma) and 10 µg/ml Alexa Fluor 546 fibrinogen conjugate (Invitrogen) in 100mM Sodium Bicarbonate buffer, pH=8.4, for 30 min then washed in 100mM Sodium Bicarbonate buffer, pH=8.4 and finally dried. Acrylamide (8%) and bis-acrylamide solution (0.48%) (Sigma) was degassed for 30 min, mixed with passivated fluorescent beads by sonication before addition of APS and TEMED. A drop of 25 µl of this mix was sandwiched between the micropatterned coverslip and a silanised (acryl-silane) glass coverslip and let to polymerize for 30 min. Gel was allowed to swell in 100mM sodium bicarbonate buffer and gently removed. Coverslip were rinced with PBS before cell plating.

The Young-modulus of the gels was estimated around 40kPa given the relative amounts of acrylamide and bis-acrylamide (Tse and Engler, 2010).

### Bead Passivation

50µl fluorescent beads (Fluorosphere #8810, Invitrogen) are incubated in 1 ml PLL-Peg (0.1 mg.ml^−1^) for 1 H at 4°C. Beads were washed 3 times in 10 mM HEPES pH 7.4 and resuspended in 150 µl washing buffer. 10 to 15 µl were added to the acrylamide gel before polymerization.

### Imaging

Live microscopy was performed on Zeiss Cell Observer Z inverted microscopes with Hamamatsu Orca flash 4.0 cameras. Fucci nuclei in time and force measurement experiments were aquired respectively with a 40x (NA=1.2) and a 63x Plan Apo (NA=1.4) objectives.

Nuclei normalized colors, Rn and Gn, were obtained by measuring each fluorescence intensity in a 5µm diameter circle manually located in the brightest part of the cell nucleus; then divided by its respective background, measured from a 5µm diameter circle manually located far from the cell.

### Traction Force Microscopy

We used the ImageJ plugin and followed the procedure previously described (Martiel *et al.*, 2015). Displacement fields were obtained from bead images taken before and after removal of cells by trypsin treatment. Images were first aligned to correct for experimental drift then cropped to produce 1000 px X 1000 px images. Displacement field was calculated by particle imaging velocimetry (PIV) on the base of normalized cross-correlation following an iterative scheme. Final grid size was 1.65 µm X 1.65 µm. Erroneous vectors where discarded owing to their low correlation value and replaced by the median value of the neighbouring vectors. Traction-force field was subsequently estimated by Fourier Transform Traction Cytometry, with a regularization parameter set to 9×10^−10^. The mechanical energy was calculated by summing the dot products of displacement with the force times the grid area: 2.72 µm^2^.

## Acknowledgements

We thank the Physiology course directors Jennifer Lippincott-Schwartz, Wallace Marshall and Rob Phillips for inviting all of us to the Marine Biology Laboratory in Woods Hole. We also thank them for their great support, thoughtful advice and remarkably good and motivating intuitions. We are very grateful to Carolyn Ott, Adam Catching and Steve Wilbert who were more than helpful both day and night, during these two weeks. We thank Neil Ganem and David Pellman for providing us the RPE1-Fucci cell line.

